# Modeling and docking antibody structures with Rosetta

**DOI:** 10.1101/069930

**Authors:** Brian D. Weitzner, Jeliazko R. Jeliazkov, Sergey Lyskov, Nicholas Marze, Daisuke Kuroda, Rahel Frick, Naireeta Biswas, Jeffrey J. Gray

**Affiliations:** Department of Chemical and Biomolecular Engineering, Johns Hopkins University, Baltimore, Maryland, USA.; T.C. Jenkins Department of Biophysics, Johns Hopkins University, Baltimore, Maryland, USA.; Department of Analytical and Physical Chemistry, Showa University School of Pharmacy, Tokyo 142-8555, Japan.; Centre for Immune Regulation, Department of Biosciences, University of Oslo, Oslo, Norway; Centre for Immune Regulation, Department of Immunology, Oslo University Hostpital Rikshospitalet, University of Oslo, Oslo, Norway.; Program in Molecular Biophysics, Johns Hopkins University, Baltimore, Maryland, USA.; Institute for NanoBioTechnology, Johns Hopkins University, Baltimore, Maryland, USA.; Sidney Kimmel Comprehensive Cancer Center, Johns Hopkins School of Medicine, Baltimore, Maryland, USA.

## Abstract

We describe Rosetta-based computational protocols for predicting the three-dimensional structure of an antibody from sequence and then docking the antibody–protein-antigen complexes. Antibody modeling leverages canonical loop conformations to graft large segments from experimentally-determined structures as well as (1) energetic calculations to minimize loops, (2) docking methodology to refine the V_L_–V_H_ relative orientation, and (3) *de novo* prediction of the elusive complementarity determining region (CDR) H3 loop. To alleviate model uncertainty, antibody–antigen docking resamples CDR loop conformations and can use multiple models to represent an ensemble of conformations for the antibody, the antigen or both. These protocols can be run fully-automated via the ROSIE web server or manually on a computer with user control of individual steps. For best results, the protocol requires roughly 2,500 CPU-hours for antibody modeling and 250 CPU-hours for antibody–antigen docking. Tasks can be completed in under a day by using public supercomputers.

## INTRODUCTION

The vertebrate adaptive immune system is capable of promoting cells to degranulate or phagocytose nearly any foreign pathogen by producing immunoglobulin G (IgG) proteins (antibodies) that recognize a specific region (epitope) of a pathogenic molecule (antigen). The ability to bind diverse antigens requires a diverse population of antibodies, which is achieved through complex processes in bone marrow and lymphatic tissues, namely V(D)J recombination and somatic hypermutation. The diversity of antibodies is astonishing; the size of the theoretical naïve antibody repertoire is estimated to be > 10^13^ in humans^1^. In addition to their biological importance, antibodies are routinely used in biotechnology as probes and diagnostics, and there are dozens of antibodies approved as therapeutics^2^.

Next-generation sequencing techniques have enabled rapid determination of large numbers of antibody sequences^1^. A limitation of these approaches is that no information about the specific atomic contacts between the antibody and antigen can be gleaned from these data sets. Atomic detail is required to consider specific antibody–antigen interactions, for example, in order to develop therapeutic antibodies or vaccines that are mimetics of extremely infectious antigens^3^. Although there are experimental methods capable of generating structural models in atomic detail (X-ray crystallography, NMR, neutron diffraction, cryo-EM), not all protein structures can be determined with these methods, and limited resources make it impossible to determine the structures of all of the sequences identified in high-throughput sequencing experiments. To bridge the sequence–structure gap, one must employ computational structure prediction methods. Perhaps more importantly, structure prediction methods are useful in diagnostics and drug discovery to define epitopes and help infer biological or therapeutic mechanisms.

The function of an antibody arises from its three-dimensional structure. The IgG isoform, the most common type of naturally occurring antibodies, consists of two identical sets of heavy and light chains arranged into a “Y” shape, with the four polypeptide chains joined by disulfide linkages. The heavy chain contains four domains, three adjacent constant domains (C_H_1, C_H_2, C_H_3) and one variable domain (V_H_), and the light chain consists of a single constant domain (C_L_) and a variable domain (V_L_). The C_H_1 and V_H_ domains interact with the C_L_ and V_L_ domains to form the antigen-binding fragment (F_ab_) or the “arms” of the Y. Within the F_ab_, both variable domains are directed away from the remaining heavy chain constant domains and make up the variable fragment (F_V_). At the tip of the FV are three complementarity determining region (CDR) loops on each chain (CDR L1–3 and CDR H1–3) that form the region of the antibody, called the paratope, that recognizes its target. This F_v_ structure is common to other antibody isoforms (IgA, IgE, etc.).

### Antibody homology modeling

The F_V_ is the focal point of the recombination and hypermutation events; as such, the primary difference among antibodies is the conformation, structural context, and chemical identity of their CDR loops. For this reason, antibody structure prediction methods focus on modeling the FV. The F_V_ can be split into two regions: framework regions, and CDR loops. The framework regions have a high degree of structural conservation, making it possible to generate accurate models of framework regions from template structures.

Similarly, analysis of antibody crystal structures has revealed that five of the six CDR loops (CDR L1–3, H1, H2) adopt a limited number of distinct structures, referred to as canonical loop conformations^4^. The canonical conformation of a particular CDR loop can typically be identified from its length and sequence. Like the framework regions, the CDRs L1–3, H1, and H2 are also modeled using template structures.

The remaining CDR loop, H3, does not adopt canonical conformations and must be modeled *de novo*. Additionally, the H3 loop lies at the interface of the two domains (V_H_ and V_L_) and can interact with residues on either chain. To account for these interactions as well as the overall geometry of the paratope, the V_L_–V_H_ orientation is optimized during H3 modeling. Accurately modeling CDR H3 and the V_L_–V_H_ orientation are typically the most challenging aspects of antibody structure prediction.

### Protein–protein docking

While accurate predictions of unbound antibody structures are informative, they are void of an important biological context: the antibody–antigen (Ab–Ag) interaction. High-resolution structures of Ab–Ag complexes give insight to the molecular mechanism by which antibodies function, a necessity for rational design of vaccines or antibody therapeutics. Structures of Ab–Ag complexes can be determined through experimental methods, however, just as with unbound antibodies, these methods are limited by their throughput and expense and are not viable for all proteins. When experimental methods cannot be used to determine complex structures, computational protein–protein interface prediction (docking) provides an alternative approach.

In general, computational docking approaches strive to sample all possible interactions between two proteins to discern the biologically-relevant interaction. Predicting a protein–protein interaction *de novo* is challenging due to the sheer number of possible docked conformations. However, the sample space can be made tractable with information about the interaction. In the case of Ab–Ag interactions, the search space is limited because the antibody paratope, comprised of the six CDR loops, is the binding site for the cognate antigen epitope.

The Rosetta SnugDock algorithm leverages the information about the flexible and/or uncertain regions of the antibody to perform robust Ab–Ag docking^5^. SnugDock simulates the induced-fit mechanism through simultaneous optimization of several degrees of freedom. It performs rigid-body docking of the multi-body (V_L_–V_H_)–Ag complex, as well as re-modeling of the CDR H2 and H3 loops, the latter of which typically contributes a plurality of atomic contacts to the Ab–Ag interaction. SnugDock can also simulate conformer selection by swapping either the antibody or the antigen with another member of a pre-generated structural ensemble. Because SnugDock samples most of the conformation space available to antibody paratopes, it can refine antibody homology models with inaccuracies in the difficult-to-predict V_L_–V_H_ orientation and CDR H3 loop.

When docking homology models, it is best if there is experimental evidence to suggest the general location of the epitope (within ~10 Å, approximately the correct side of the antigen domain), and in this protocol paper, we describe the local docking procedure in detail. If no information is available about the epitope, there are several programs that perform global docking or epitope prediction^6^. In particular, there are two fast-Fourier transform (FFT) rigid-body docking approaches that implement antibody-specific energy potentials: PIPER^7^ with the antibody-ADARS potential^8^, and ZDOCK^9^ with the Antibody i-Patch potential^10^. FFT rigid-body approaches are fast, but they cannot account for antibody motions upon antigen binding or compensate for errors in the initial homology model; SnugDock is the only flexible-backbone antibody docking method. It can provide a global-antigen docking alternative but it is slower and, like others, can produce false-positive epitope predictions^5^. For local docking, SnugDock has been demonstrated to produce high-quality models when using an antibody homology model or crystal structure and the unbound antigen crystal structure as input^5^. In addition, SnugDock approaches used in the CAPRI blind docking challenge^11^ produced the best structure among all predictors for a flexible-loop target. CAPRI uses star-based rankings (*** = high quality, ** = medium, * = acceptable, 0 = incorrect)^12^. Examining the highest attained CAPRI quality among the ten lowest-scoring docked models (starting with a homology modeled antibody), SnugDock currently produces 1***, 10**, and 4* models in a test set of 15 antibody-antigen targets (Table 1). These performance data are improved since the original SnugDock publication^5^ due to updates in the energy function^13^ and a switch to the kinematic loop closure (KIC) loop modeling method.^14-16^

**Table 1.** Local Ab–Ag docking benchmark results. Co-crystal PDB IDs indicate the native complex. PDB IDs listed under the “Type” column also indicate the use of unbound (U) or bound (B) component structures (as available). Model quality is defined by the CAPRI ranking criteria represented by a number of stars or a zero (0). Three, two, and one star(s) indicate high, medium, and acceptable quality, respectively, and a zero indicates incorrect models. For the “Rigid-Body” and “SnugDock” columns, the quality of the lowest-scoring model, by interface energy, is reported (an “f” indicates a strong energy funnel defined as five or more of the ten lowest-scoring models being medium quality or better). Ensemble SnugDock simulations were run with multi-template grafting and a CDR H3 kink constraint. CAPRI Summary lines summarize model quality for all targets. CAPRI Summary Top 10 takes the highest-quality model from the ten lowest-scoring models.

| Co-crystal<br>(PDB ID) | Type (Ab-Ag) | CDR H3<br>Length | Rigid-Body<br>Dock Xtal | Rigid-Body<br>Dock Model | SnugDock | Ensemble<br>SnugDock |
| --- | --- | --- | --- | --- | --- | --- |
| 1mlc | U(1mlb)-U(1lza) | 7 | 0 | 0 | ** | * |
| 1ahw | U(1fgn)-U(1boy) | 8 | 0 | * | * | ** |
| 1jps | U(1jpt)-U(1tfh) | 8 | ** | * | * | ** |
| 1wej | U(1qbl)-U(1hrc) | 8 | 0 | 0 | 0 | * |
| 1vfb | U(1vfa)-U(8lyz) | 8 | * | * | 0 | * |
| 1bql | B-U(1dkj) | 7 | *** f | 0 | * | 0 |
| 1k4c | B-U(1jvm) | 9 | ** f | 0 | * | ** |
| 2jel | B-U(1poh) | 9 | *** f | 0 | ** | * |
| 1jhl | B-U(1ghl) | 9 | *** f | 0 | 0 | ** |
| 1nca | B-U(7nn9) | 11 | *** f | 0 | 0 | * |
| 2bdn | B-B | 8 | *** f | * | * | * |
| 1ynt | B-B | 9 | *** f | *** f | ** f | *** f |
| 2aep | B-B | 9 | *** f | ** | * | 0 |
| 2b2x | B-B | 10 | *** f | * | 0 | ** |
| 1ztx | B-B | 10 | *** f | * | ** f | 0 |
| CAPRI Summary Top Decoy |  |  | 9***/2**/1* | 1***/1**/6* | 0***/4**/6* | 1***/5**/6* |
| CAPRI Summary Top 10 Decoys |  |  | 9***/4**/2* | 1***/6**/8* | 1***/10**/4* | 2***/11**/2* |
| No. of Funnels |  |  | 10 | 1 | 2 | 1 |

### Protocol overview

The protocol described in this paper enables a user to generate a structural model of an antibody from its sequence and a structural model of an antibody–antigen complex from structures of the antibody and its antigen (Fig. 1).

**Figure 1.**
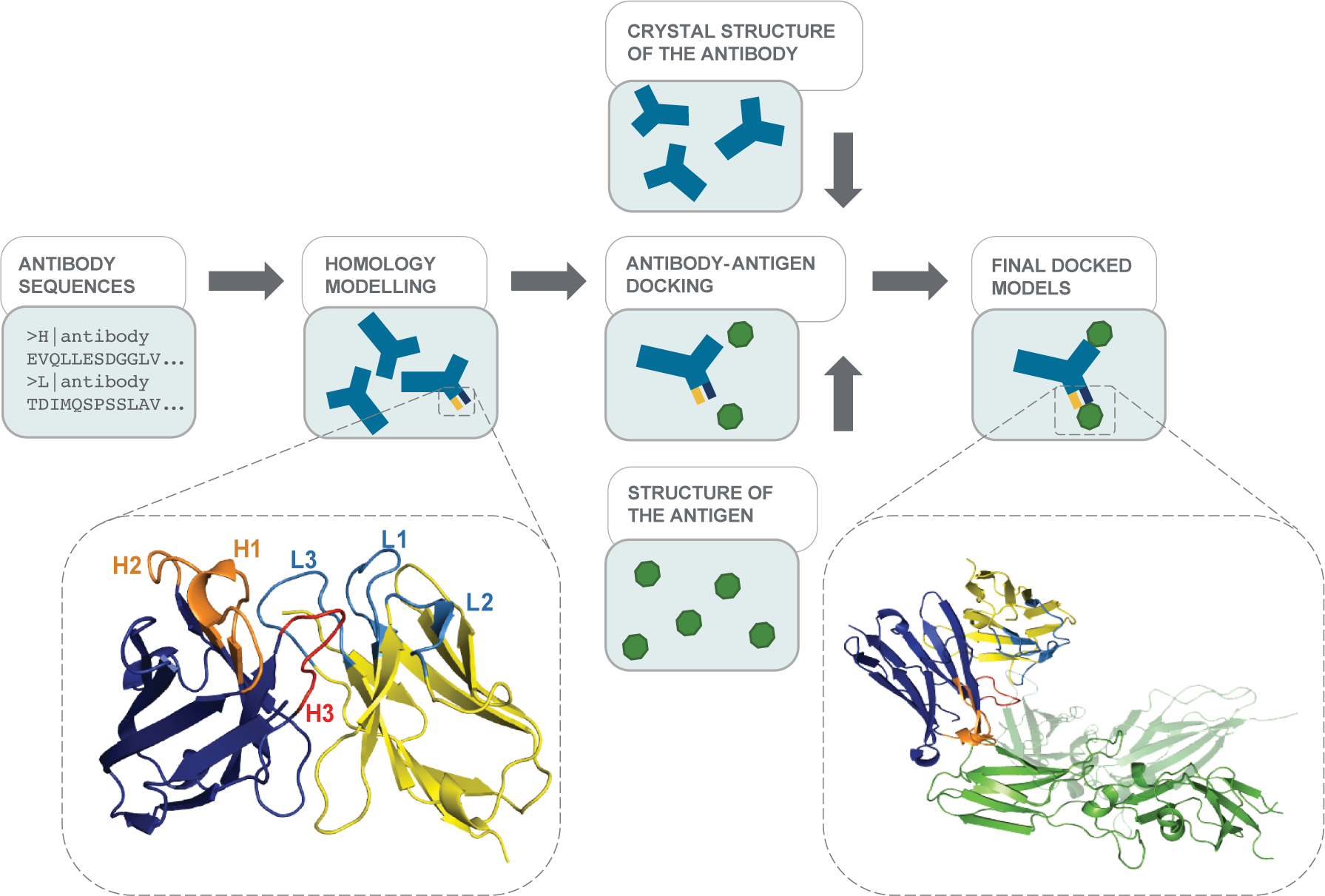
A schematic of the modeling protocols. The structure on the left shows the F_V_ antibody domains predicted by homology modeling (heavy chain in dark blue with CDR H1 and H2 loops in orange and CDR H3 loop in red; light chain in yellow with its CDR loops in light blue). The structure on the right depicts an antibody–antigen structure output by docking (antigen in green).

**Figure 2:**
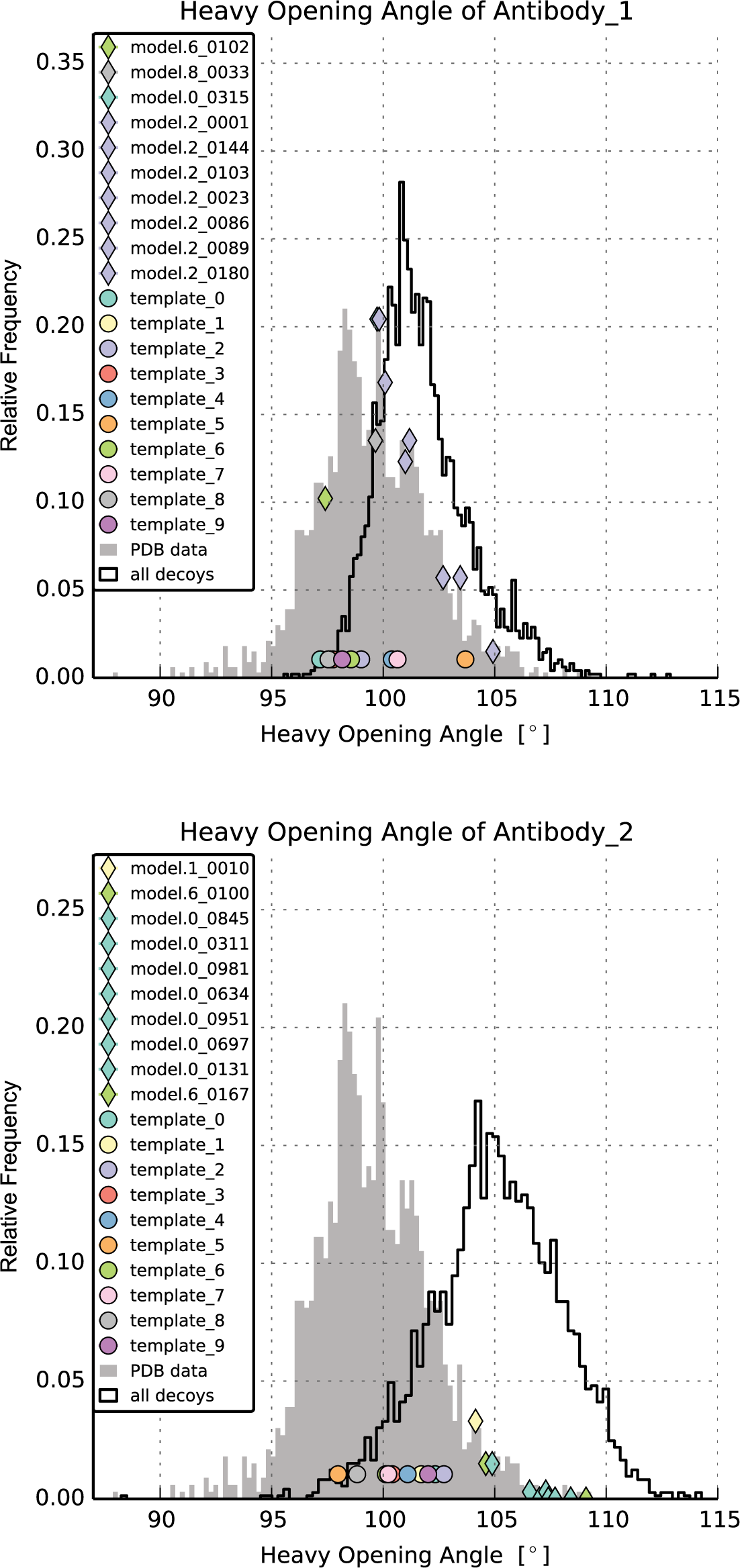
Example output of plot_LHOC.py. The two plots show distributions of the Heavy Opening Angle^18^ as obtained by plot_LHOC.py for two different antibodies. The 10 distinct light-heavy orientation templates are represented by the circles. The ten top-scoring models after H3 loop modeling are represented by the diamonds with the fill color corresponding to the starting template; in the legend, these points are ordered from smallest to largest metric value. For Antibody_1, the angles sampled by Rosetta overlap with the angles observed in antibody crystal structures. The ten top-scoring models are close to the center of the distribution. In Antibody_2, most of the angles sampled are found rarely or not at all in antibody crystal structures. The ten top-scoring models are also shifted to larger angles than typically found in antibodies. For Antibody_2, the user might consider trying alternate light-heavy orientation templates (Step 10).

### Protocol overview: Antibody homology modeling (steps 1-8)

Generating a structural model of an antibody from sequence in RosettaAntibody uses homology modeling techniques, that is, it uses segments from known structures with similar sequences. As described in detail below, the input sequence is split into several components. For each component, RosettaAntibody searches a curated database of known structures for the closest match by sequence and then assembles those structural segments into a model. That model is then used as the input for the next stage in which the CDR H3 loop is modeled and the V_L_–V_H_ orientation is optimized.

### Numbering the residues in the sequence

The RosettaAntibody protocol identifies the CDRs of the input antibody sequence through regular expression matching to the Kabat CDR definition^17^, and it numbers the antibody residues according to the Chothia scheme^4^.

### Template selection

For each structural component considered (FRL, FRH, CDRs L1–3, H1–3), templates are selected by maximum sequence similarity using a BLAST-based method with custom databases constructed from high-quality structures in the PDB. Canonical CDR conformations are based on length, so we use separate databases for each loop–length combination. For example, ten-residue H1 loops and eleven-residue H1 loops are separate BLAST-formatted databases.

The results for each structural component are sorted by BLAST bit score, and the sequence with best score is selected as the template.

### Initial V_H_-V_L_ orientations

The initial V_L_–V_H_ orientation is selected in much the same way, with the exception that ten V_L_–V_H_ templates are selected rather than a single one^18^. Starting from the list of all possible templates ordered by bit score, the best match is selected as the first template. To diversify the initial V_L_–V_H_ orientations, all templates with similar V_L_–V_H_ orientations (0.5 OCD, see Marze & Gray^18^) to this template are pruned from the list. The best match remaining in the list is selected as the second template, and candidate templates similar to the second template are now removed from the list. This winnowing is repeated to create ten distinct templates. One grafted model will be created from each of these ten initial V_L_–V_H_ orientations.

### Grafting CDR templates

Once the initial V_L_–V_H_ orientations are set, the CDR templates are grafted onto each framework region by superposing the two overlapping residues on either side of the loop with their corresponding residues on the framework regions. The graft points are then adjusted using Cyclic Coordinate Descent (CCD)^19,20^ to prevent unphysical bond lengths and angles from being incorporated into the model. Finally, the structure is relaxed^21,22^ via iterations of side-chain optimization and gradient-based minimization while constraining the backbone and side-chain heavy atoms to find a native-like conformation at a local energy minimum in Rosetta’s score function.

### All-atom refinement of CDR H3 and the V_L_–V_H_ orientation

The grafted models are crude and must be refined, particularly in the CDR H3 loop and the V_L_–V_H_ orientation. The H3 loop is first completely remodeled in the context of the antibody framework using the next-generation KIC (NGK) loop modeling protocol^16^. For speed, the H3 loop side chains are each reduced to a single low-resolution pseudo-atom, and to ensure sampling of the C-terminal kink conformation, atomic constraints are applied to the governing score function^23^. For subsequent high-resolution refinement, the all-atom CDR H3 side chains are recovered, all CDR side chains are repacked, and the CDR side chains and backbones are minimized. The V_L_ and the V_H_ domains are re-docked with a rigid-backbone RosettaDock protocol24,25 to remove any clashes created by the new H3 conformation, and the antibody side chains are again repacked. Using NGK, H3 is refined again in the context of the updated V_L_–V_H_ orientation. The CDRs are packed and minimized again, and the model is saved as a candidate structure, or decoy. The first grafted model is used as the starting point for 1,000 refined models, or decoys, and the other grafted models are each used as the starting point for 200 decoys, for a total of 2,800 decoys. The decoys are sorted by Rosetta score, and the lowest-scoring ones are given as the final models.

### Protocol overview: antibody–antigen docking (steps 11-16)

Computational docking can be used to generate models of Ab–Ag complexes. In general, docking entails (1) roughly identifying (within 8 Å) the interacting interface through either experiment or global docking and (2) refining the initial model through local docking. Below we describe local docking with SnugDock in detail.

### Generating the starting model

SnugDock requires, as an input, a putative Ab–Ag complex that contains a reasonable interface^26^. The complex can be composed of single structures or sets of structures (ensembles, see Box 1). The interface defines the local search, between the antibody CDRs and the antigen. Initial models are often based on experimental results that identify interacting residues at the Ab–Ag interface, such as mutagenesis or chemical crosslinking assays. In the absence of experimental results, a global docking approach such as ZDOCK/iPatch^10^ or PIPER/ADARS^8^ can generate putative complexes for refinement. Global docking can also be achieved with SnugDock, albeit at a higher computational expense.

##### Box 1 | Increasing sampling during docking by incorporating backbone structural ensembles

In Rosetta, an ensemble is a set of discrete conformations of a protein structure. SnugDock uses ensembles to approximate backbone conformational flexibility by sampling conformations from the ensemble during docking. Through this approach, not only does the protocol explore more conformational space than standard docking, but it can also compensate for model error, for example by using an ensemble of models produced by a modeling approach in a previous step such as RosettaAntibody.

Rosetta ensembles can be converted directly from NMR ensembles, or they can be generated using any method that induces structural diversity, such as molecular dynamics or various Rosetta refinement protocols. The ensembles typically span small structural variations of 1-2 Å backbone RMSD^25^. Rosetta’s relax protocol (unconstrained)^21,22^ or KIC protocol^14-16^ are suggested to generate docking ensembles for antigens. In addition, RosettaAntibody creates ensembles of antibodies by default. More on how to generating and docking ensembles can be found in Chaudury and Gray^25^ and in Rosetta’s documentation^37^.

Antigen or antibody structures that have not been generated by a Rosetta protocol need to be refined before being placed in contact. Refinement, commonly referred to as the Relax protocol^21,22^, entails iterations of side-chain optimization and gradient-based minimization in Rosetta’s score function. The Relax protocol samples local conformational space around the starting structure to identify an energetic minimum in the score function. Through this process, Rosetta-identified non-idealities (such as van der Waals bumps) are abated. Once the partners have been refined, a putative complex can be assembled and prepacked. Prepacking optimizes side-chain conformations to prevent biasing toward the input complex model’s side-chain conformations, ensuring uniform scoring of all potential bound complex states.

### Performing docking

SnugDock iteratively performs multi-body docking of both the Ab–Ag and V_L_–V_H_ orientations and remodeling of the H2 and H3 CDR loops. Prior to docking, the prepacked starting Ab–Ag complex is subject to three rigid-body perturbations: (1) a randomized rotation about the Ab–Ag primary axis, (2) a small-magnitude random translation, and (3) a small-magnitude random rotation. Docking operates in two phases: low-resolution mode, where side chains are represented by a single pseudoatom located at the centroid of the side-chain heavy atoms, and high-resolution mode, where all protein atoms are explicit. Low-resolution mode consists of two types of interspersed Monte Carlo moves: rigid-body Ab–Ag translation and rotation, and backbone ensemble conformer swaps. Additionally, at the end of low-resolution mode, the H2 & H3 loops are refined. High-resolution mode consists of a 50-step Monte Carlo trajectory where each move is selected from a set of five possible moves: rigid body Ab–Ag docking (40%), rigid body V_L_–V_H_ docking (40%), CDR minimization (10%), H2 loop refinement (5%), and H3 loop refinement (5%), where the percentages indicate the probabilities of selecting each move. Each trajectory results in one decoy. Typically, SnugDock is used to generate a total of 1,000 decoys, with the low-scoring decoys most likely to be near the native conformation.

### Incorporating experimental data into the simulation

Two main types of experimental data that inform the Ab–Ag binding mode can be incorporated into SnugDock. First, knowledge about specific residues or pairs of residues that interact across the interface can be used to guide docking. This information could, for example, be derived from alanine scanning or other mutagenesis experiments. Second, knowledge about the epitope and the overall Ab–Ag orientation can be incorporated. Binding patch data may be derived from different experiments, including hydrogen/deuterium exchange or chemical crosslinking of the binding partners with subsequent analysis by mass spectrometry. Other methods for epitope mapping may also be suitable.

Depending on the type of experimental data available, there are different ways of incorporating it into the docking simulation. High-confidence residue–residue interactions can be preserved with the use of atom pair constraints. Less-specific and poorly-characterized interactions (hydrophobic pockets, ambiguous H-bonds) can be loosely constrained with ambiguous and site constraints. Predicted epitopes and binding patches can be sampled by properly placing the SnugDock input structure and adjusting the size of the initial starting move. For further information on incorporating experimental constriants, see the Rosetta documentation^27^.

### Caveats, challenges and pitfalls

There are several caveats associated with computational modeling of antibodies and docking of antibodies and antigens. Keeping these caveats in mind, the user should critically assess each prediction (see Box 2). RosettaAntibody is a homology modeling approach and can be hampered by template availability. For example, challenging targets include heavily engineered antibodies or antibodies derived from a species that diversifies its antibodies through gene conversion, such as chickens or rabbits. Errors in the FR and CDR L1–3, H1, H2 loops are typically small (no greater than 1 Å backbone RMSD to native)^28^. The V_L_–V_H_ orientation, correctly captured by RosettaAntibody in 43 of 46 benchmark antibody targets18. The CDR H3 loop, on the other hand, is modeled *de novo*, and loop model quality decreases with loop length. In the KIC loop benchmark,^16,29^ loops of 12–17 residues are modeled to near 1 Å backbone RMSD relative to the native structure–the average human CDR H3 falls within that range with an average length of 15 residues (IMGT definition)^30^. However, the benchmark is measured by modeling loops on crystallographic frameworks, whereas in a blind context, CDR H3 loops are modeled on a homology frameworks, which introduces uncertainty in the loop environment. Nevertheless, in a recent assessment^23^ Rosetta Antibody produced models with CDR H3 loops within 1.59 Å backbone RMSD to native and sub-angstrom accuracy in all other regions.

##### Box 2 | Assessing antibody modeling and antibody–antigen docking results

The user must critically analyze computational models. Models output by Rosetta should be ranked according to score. In most simulations, approximately 90% of the models will be non-native-like, and they will occupy a bulk score range of 30-50 Rosetta Energy Units (REU). Only about 1-5% of models will be native-like, and they will have scores ranging from within the bulk score range to a well of 5-10 REU below the bulk score range. If a cluster of structurally related models lie in the well below the bulk score range, these are likely native-like, with deeper scoring and more populated wells providing higher confidence.

###### Antibody model assessment

First, assess the physical feasibility of the lowest-scoring models by eye in a molecular visualization package such as PyMOL. It is important to check for obvious flaws that can occur in rare circumstances such as polypeptide chain breaks or backbone clashes, particularly within the CDRs and at their graft points. The accuracy of the non-H3 CDR loops should be further assessed by comparing the CDR cluster of the grafted loop with the cluster of the input sequence as identified by North et al.^36^ (see step 5). Next, ensure the components of the V_L_–V_H_ orientation lie within nature’s distribution (step 9), as models with outlying orientations are not likely to be native-like; an exception to this rule can be made if the V_L_–V_H_ orientation grafting templates and Rosetta sampling all lie far toward the edge of nature’s distribution.

###### Ab–Ag docking model assessment

Make sure that the lowest-scoring models make good contacts between the antigen and the antibody paratope. Higher confidence can be assigned to models with large (~1200 Å), complementary interfaces,^26^ as well as those in which the H3 CDR loop makes several specific contacts. If experimental data suggest an antigen binding site, ensure the paratope contacts at this site.

While SnugDock explicitly samples the CDR H2 or H3 loop conformation and V_L_–V_H_ orientation to account for model uncertainty introduced during homology modeling and to sample the regions most likely to undergo conformational changes upon antigen binding, it does not explicitly sample backbone degrees of freedom of the antigen or of non-CDR-H2 or H3 regions of the antibody. Thus, if the unbound and bound conformations differ substantially or if the homology models are poor, it could be difficult or impossible to model the docked complex
accurately^31^. Despite this complication, SnugDock has successfully predicted Ab–Ag complexes from homology models^5^.

### Availability

RosettaAntibody and SnugDock can be run via a public webserver (http://rosie.rosettacommons.org), python bindings (PyRosetta, http://www.pyrosetta.org) and through local installations of Rosetta. Rosetta is distributed as source code and licenses are available from the RosettaCommons (http://www.rosettacommons.org) free of charge for academic and non-profit users. Rosetta can be installed on UNIX-like operating systems (including Mac OS X).

## MATERIALS

### EQUIPMENT

#### Homology modeling data

- Primary amino-acid sequence of the variable domain of the light and heavy chains.

#### Docking data

- PDB-formatted file of the antigen structure.
- PDB-formatted file of the antibody structure, from the homology modeling output.
- Both of these can be single structures or an ensemble of structures.

#### Software for running simulations via ROSIE web server

- Modern web browser

#### Hardware for running simulations manually (optional)

- Workstation with multi-core CPU(s) running a POSIX compliant operating system (*e.g.*, GNU/Linux, OS X)

OR

- a Linux-based cluster. Several public facilities are available. For example, the U.S. National Science Foundation’s provides clusters like Stampede through the Extreme Science and Engineering Discovery Environment (XSEDE, www.xsede.org). In Europe, the Partnership for Advanced Computing in Europe (PRACE, www.prace-ri.eu) provides access to clusters like JUQUEEN. Resources like the Norwegian Metacenter for Computational Science (Notur, www.notur.no) or Japan’s supercomputer facilities of National Institute of Genetics (sc.ddbj.nig.ac.jp) and of Human Genome Center at the University of Tokyo (hgc.jp) are also suitable.

#### Software for running simulations locally (optional)

- The Rosetta software suite, available at www.rosettacommons.org/software
  - Compilation instructions available at www.rosettacommons.org/build. **? TROUBLESHOOTING**
  - Support for any issues encountered that are not covered in this manuscript can be addressed on the Rosetta user forums: www.rosettacommons.org/forum
- BLAST+ (version 2.2.28 or later), available at ftp://ftp.ncbi.nlm.nih.gov/blast/executables/blast+/LATEST/
- Text editor (e.g., vim, emacs, nano)
- Optional: Python (www.python.org) or R (www.r-project.org) for analyzing results
- Optional: A molecular visualization package for viewing results and customizing starting structures for docking. Recommended packages include PyMOL (www.pymol.org)^32^, UCSF Chimera (www.cgl.ucsf.edu/chimera)^33^, and Kinemage (kinemage.biochem.duke.edu)^34^

## PROCEDURE

The simplest way to create antibody and antibody–antigen complex structures is through the use of the ROSIE web server (rosie.rosettacommons.org)^35^. On ROSIE, the Antibody app uses the input antibody sequence to generate a homology model, and the SnugDock app uses the antibody model(s) and an antigen structures for docking. Both operations are entirely automated with a minimum of user input.

For greater control of the operation, we describe below the steps to run the protocols manually, including the key points for checking intermediate data and intervening with alternate choices. Users with structures of the unbound antibody and antigen can skip to docking stage (step 11).

### I. Antibody Homology Modeling

#### Construction of a grafted F_v_ model TIMING 75-90 minutes

1. Set up your terminal. After installing BLAST+ and Rosetta (see Materials), launch an interactive terminal (*e.g., Terminal* on mac or *xterm* on Linux) and set path variables to the executable programs needed as follows (bash syntax):

~~~
export ROSETTA=~/Rosetta
export ROSETTA3_DB=$ROSETTA/main/database
export ROSETTA_BIN=$ROSETTA/main/source/bin
export PATH=$PATH:$ROSETTA_BIN
~~~

In the first line above, replace “~” with the parent directory where you installed Rosetta on your machine. Similarly, be sure the PATH variable includes the blastp program (e.g. `export PATH=$PATH:/path/to/blastp` where `/path/to/blastp` is replaced with the directory containing the *blastp* executable. These path settings may be added to a configuration file such as `.bashrc` so they are automatically set each time a terminal is open (logged into).
2. Create a working directory and navigate to it:

~~~
mkdir /path/to/my_dir
cd /path/to/my_dir
~~~
3. Obtain the amino acid sequences for the variable domain of your antibody (light chain and heavy chain) and save them in FASTA format (in your working directory) with the heavy and light chains noted in the comment lines, as follows:

~~~
> heavy
VKLEESGGGLVQPGGSMKLSCATSGFRFADYWMDWVRQSPEKGLEWVAEIRNKANNHATYYAESVKGRFTI
SRDDSKRRVYLQMNTLRAEDTGIYYCTLIAYBYPWFAYWGQGTLVTVS

> light
DVVMTQTPLSLPVSLGNQASISCRSSQSLVHSNGNTYLHWYLQKPGQSPKLLIYKVSNRFSGVPDRFSGSG
SGTDFTLKISRVEAEDLGVYFCSQSTHVPFTFGSGTKLEIKR
~~~
4. Use Rosetta’s grafting application to find suitable templates and graft them together to obtain a crude model of the antibody. Execute the application with the line below.

~~~
antibody.macosclangrelease \
      -fasta antibody_chains.fasta | tee grafting.log
~~~

The application will output a directory called `grafting`. The PDB-formatted files named `model-0.relaxed.pdb, model-1.relaxed.pdb, …, model-9.relaxed.pdb` will be your input for the H3 modeling. The “`| tee grafting.log`” part of the command records all the program output in the file `grafting.log` for later review. The “\” permits the command to be spread across multiple lines rather than just one. **? TROUBLESHOOTING**

#### (Optional) Check grafted template structures TIMING: 10 minutes – 2 hours

5. Assign the CDR loops in your models to the CDR loop clusters described by North et al.^36^ and check whether the chosen templates are suitable. Run the cluster identification application as follows:

~~~
identify_cdr_clusters.macosclangrelease \
      -s grafting/model-*.relaxed.pdb \
      -out:file:score_only north_clusters.log
~~~

North *et al*. clustered all CDR loop structures by their backbone dihedral angles and named them by CDR type, loop length and cluster size (e.g. “H1-13-10” is the 10^th^ most common conformation for 13-residue H1 loops). Occasionally, Rosetta chooses templates that are rare or inconsistent with the sequence preferences observed by North *et al*. For example, if Rosetta recommends the H1-13-10 cluster, the user might
also consider the H1-13-1 cluster. Tables 3-7 of North *et al*. present consensus sequences for each cluster that can inform this decision. Loops and clusters with proline residues are also worth a manual examination. Several clusters of North *et al*. are contingent on the presence of prolines in particular locations (e.g. L3-9-*cis7*-1 has a *cis*-proline at position 7). Because RosettaAntibody relies on BLAST to choose loop templates, occasionally a loop from an uncommon non-*cis*-proline cluster (e.g. L3-9-2) is chosen. In such cases it is best to manually select a loop template from the well-populated *cis*-proline cluster.
6. If desired, rerun grafting to replace a template with one from a manually-specified source structure. Use the antibody command line as above with an extra flag to specify a template. Follow the below example: to force Rosetta to use the CDR H1 loop from the PDB 1RZI as the template in the model, add the flag `-antibody:hi_tempiate lrzi`. Select templates for other regions accordingly:

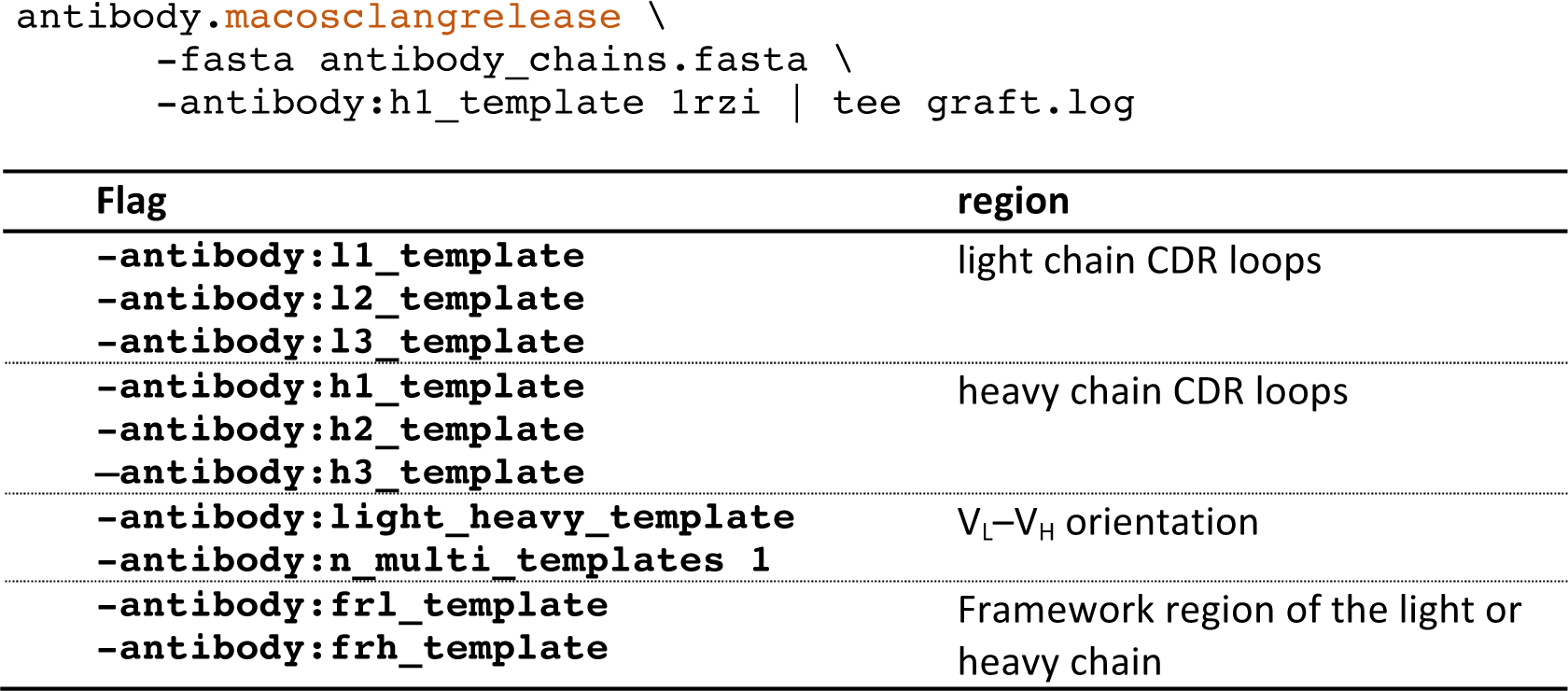

#### H3 modeling TIMING 1 hour to 4 days

7. Copy the set of standard H3 modeling flags to your working directory and create a directory for the H3 modeling output:

~~~
cp $ROSETTA/tools/antibody/abH3.flags .
mkdir H3_modeling
~~~
8. Run Rosetta’s `antibody_H3` application on the 10 models generated during grafting. This step requires 2,500 CPU hours and is often performed in parallel on a computer cluster (see Box 3).

##### Box 3 | Using Rosetta on different platforms and running in parallel

**Rosetta on different platforms.** Throughout this protocol executables are suffixed by the platform and mode for which they were compiled (i.e. `antibody.macosclangrelease` indicates that the antibody executable was compiled on a `MacOS` operating system using the Clang compiler and it was compiled in release mode). The suffix is highlighted in orange throughout (`.macosclangrelease`). On other platforms you will replace this string with your operating system and compiler (for example, GNU/Linux platforms with gcc as the compiler will default to `.linuxgccrelease`). Additionally, the suffix is prefixed by `.mpi` (`.mpi. linuxgccrelease`) when the executable is built for the message passing interface (MPI) by an MPI compiler. MPI-compatible executables can communicate with one another for parallel processing, and some Rosetta executables use MPI non-trivially. However, most standard Rosetta applications are trivially parallelizable (“embarrassingly parallel”) and thus capable of running on both MPI and non-MPI systems.

**Running in parallel.** An example of how to locally run a non-MPI executable in parallel is given in step 8. In general, add the `-multiple_processes_writing_to_one_directory` flag to your command line, and then execute multiple instances of the process. This procedure works on a single desktop computer with multiple CPUs or remotely on a supercomputer cluster. However, running a Rosetta executable on a cluster strongly depends on the hardware configuration and available software (e.g. workload management software).

For example, to run a non-MPI executable via HTCondor: (1) save the standard command line as an executable bash script, (2) write a submit description file specifying the executable bash script and the number of processes to execute, and (3) use the `condor_submit` command with the description file as an argument to submit your jobs to the cluster.

On the other hand, MPI executables can be run in parallel locally by prepending the command line with the `mpirun -n XX` command, where XX is the number of processes to run, if your machine is configured to use the Open MPI library. Again, the exact depend on the specific cluster configuration. For example, to run an MPI executable on Stampede via the *slurm* workload manager: (1) save the standard command line as an executable bash script, (2) write a *slurm* batch script specifying the executable bash script and the number of tasks, and (3) use the `sbatch` command with the bash script as an argument to submit your jobs to the cluster.

For a Mac workstation, use the following command line:

~~~
$ROSETTA_BIN/antibody_H3.macosclangrelease \
             @abH3.flags \
             -s grafting/model-0.relaxed.pdb \
             -nstruct 1000 \
             -antibody:auto_generate_kink_constraint \
             -antibody:all_atom_mode_kink_constraint \
             -multiple_processes_writing_to_one_directory \
             -out:file:scorefile H3_modeling_scores.fasc \
             -out:path:pdb H3_modeling > h3_modeling-0.log 2>&1 &
~~~

- `-s` specifies the input file (one of the grafted models generated in step 2).
- `-nstruct` specifies the number of structures generated, which should be 1000 for model.o.pdb and 200 each for all other grafted models.

The expected output is the specified number of PDB files as well as a score file named `H3_modeling_scores.fasc`. All these files will appear in an output directory named `H3_modeling/`.

To trivially run in parallel, simply repeatedly execute the above command (changing input models, number of structures, and the output log as you wish). Each time the command is executed, an *antibody_H3* process is run in the background.

!Caution: Generating the 2,800 antibody structures takes approximately 2,500 CPU hours. Running 24 processes in parallel, on a modern 24-CPU workstation, expect ~4 days of run time. Distributing the work over nodes on a supercomputer can reduce this time to hours (see Materials).

#### (Optional) Check V_L_–V_H_ orientation TIMING : 5 min

9. Check whether the V_L_–V_H_ orientations of the antibody models are close to the orientations observed in antibody crystal structures found in the PDB. Run the python script `plot_LHOC.py` using the following command line:

~~~
python
$ROSETTA/main/source/scripts/python/public/plot_V_L__V_H__orientational_coo
rdinates/plot_LHOC.py
~~~

This script will create a subfolder (`lhoc_analyis`) with separate plots for each of the four LHOC metrics. Each plot shows the native distribution of V_L_–V_H_ orientations (grey), the orientations sampled by Rosetta (black line) as well as the top 10 models (labeled diamonds) and the 10 different template structures generated during step 2 (dots). Antibody models that are outside the native distributions are unlikely to be correct.

#### Choose final antibody models TIMING 10 min

10. Choose 10 of the antibody models as an ensemble for docking. The following criteria may be useful to consider as docking with ensembles aims to increase conformational diversity and sampling:

a. Select models with the lowest total score - these are purportedly native-like
b. Select models with natural V_L_–V_H_ orientation, falling within the observed distribution (grey).
c. Select models derived from different templates to maintain diversity.

If all ten top-scoring models are outside the native distribution, consider returning to step 6 and manually select new templates for the relative orientation of the V_L_ and V_H_ chains by using the `-antibody:light_heavy_template` **flag** (***e.g.,*** `antibody.macosclangrelease -antibody:light_heavy_template 1ABC`)

### II. Antibody–Antigen Docking TIMING 1 hour

11. Prepare the antigen and antibody for docking. Format your antigen (and antibody if you are not using a homology model produced by Rosetta Antibody) PDB file so it can be read by Rosetta. Run the following script:

~~~
$ROSETTA/tools/protein_tools/scripts/clean_pdb.py antigen.pdb C
~~~

Where `antigen.pdb` is a PDB file of your antigen and C is the one-letter chain identifier(s) for the antigen chain(s) in the PDB file.

#### (Optional) Refine antibody in Rosetta’s score function TIMING 10 min

12. If you are not using an antibody model produced by Rosetta, you must refine the antibody structure by running the relax application. The command line is:

~~~
relax.macosclangrelease \
      -s antibody.pdb \
      -relax:constrain_relax_to_start_coords \
      -relax:ramp_constraints false \
      -ex1 \
      -ex2 \
      -use_input_sc \
      -flip_HNQ \
      -no_optH false
~~~

You may also wish to generate an ensemble of antibody structures, see Box 2.

#### Prepacking TIMING 10 min

13. Generate a PDB file that contains both your antibody and your antigen in the following order: light chain of your antibody (L), heavy chain of your antibody (H), and antigen (A). There are several ways to create and modify a PDB file. For example, with PyMOL:

a. Load the antibody in a PyMOL session.
  i. If it is a model from Rosetta Antibody, the chains will already be labeled as H and L. Otherwise, use the alter command to change the chain ID of a selection:

~~~
alter chain A, chain=‘H’
alter chain B, chain=‘L’
~~~
b. Load the antigen into the same PyMOL session. Change the antigen chain ID in a similar fashion. !Warning: if antigen chains share an ID with the antibody, you will have to be more specific with your selections (*e.g.*, `alter chain H and antigen, chain=‘A’`).
c. Reorient the antibody and antigen using the RotO, MovO, and MvOZ editing commands. Alternatively, one can also use the translate command (*i.e.* `translate [x,y,z], selection`). If you know an approximate binding location, adjust the orientation accordingly.
d. Save both objects in the same PDB file:

~~~
save antibody_antigen_start.pdb, chains L+H+A
~~~
14. To ensure low-energy starting side-chain conformations, prepack the monomers:

~~~
docking_prepack_protocol.macosclangrelease \
      -in:file:s antibody_antigen_start.pdb \
      -ex1 \
      -ex2 \
      -partners LH_A \
      -ensemblel antibody_ensemble.list \
      -ensemble2 antigen_ensemble.list \
      -docking:dock_rtmin
~~~

`antibody_ensemble.list` is a text file that contains filenames with absolute paths to the ten antibody models selected after antibody modeling. In the case that you have a single crystal structure, you can omit the `-ensemblel` flag. If antigen flexibility is expected, a family of structures can be created with other Rosetta applications (see Box 1). The text file `antigen_ensemble.list` will contain the filenames of your antigen (using absolute paths). NMR starting structures must be split (i.e. each model should be in its own PDB file). To use a single antigen structure, omit the `-ensemble2` flag.

#### Docking TIMING 15 min

15. Dock the antibody to the antigen. As in step 8, this is an expensive computational step and you have the option of running a single process, multiple processes on one machine, or splitting the job across processors on a supercomputer (see Box 3). Using the executable for an MPI-based computing cluster as an example, the command line for docking is:

~~~
snugdock.mpi.linuxgccrelease \
       -s prepack/antibody_antigen_start.prepack.pdb \
       -ensemblel antibody_ensemble.list \
       -ensemble2 antigen_ensemble.list \
       -antibody:auto_generate_kink_constraint \
       -antibody:all_atom_mode_kink_constraint
       -nstruct lOOO
~~~ **? TROUBLESHOOTING**

## TIMING

Here we report the time to generate a single, docked model from antibody sequence and antigen crystal structure. Typically, however, thousands of models are generated, so we indicate in parentheses the timing for the full, recommended simulations. These time estimates were computed on a 2 × 2.4 GHz Quad-Core Intel Xeon processor; timing will vary for other computer configurations.

| Step | Human Time | CPU Time per Model | Total CPU Time |
| --- | --- | --- | --- |
| (1–4) Construction of grafted Fv models | 5 min | 20 min | 200 min |
| (5) Check grafted models | 10 min | 1 min | 10 min |
| (7–8) H3 modeling | 5 min | 50 min | 2500 hrs |
| (9) Check $V_L$–$V_H$ orientation | 5 min | 1 min | 10 min |
| (10) Choose models | 10 min | 5 min | 5 min |
| (11) Prepare antibody and antigen for docking | 5 min | 15 min | 15 min |
| (12) Refine antibody in Rosetta's score function | 5 min | 20 min | 20 min |
| (13–14) Prepacking | 5 min | 10 min | 10 min |
| (15) Docking | 5 min | 15 min | 250 hrs |

## TROUBLESHOOTING

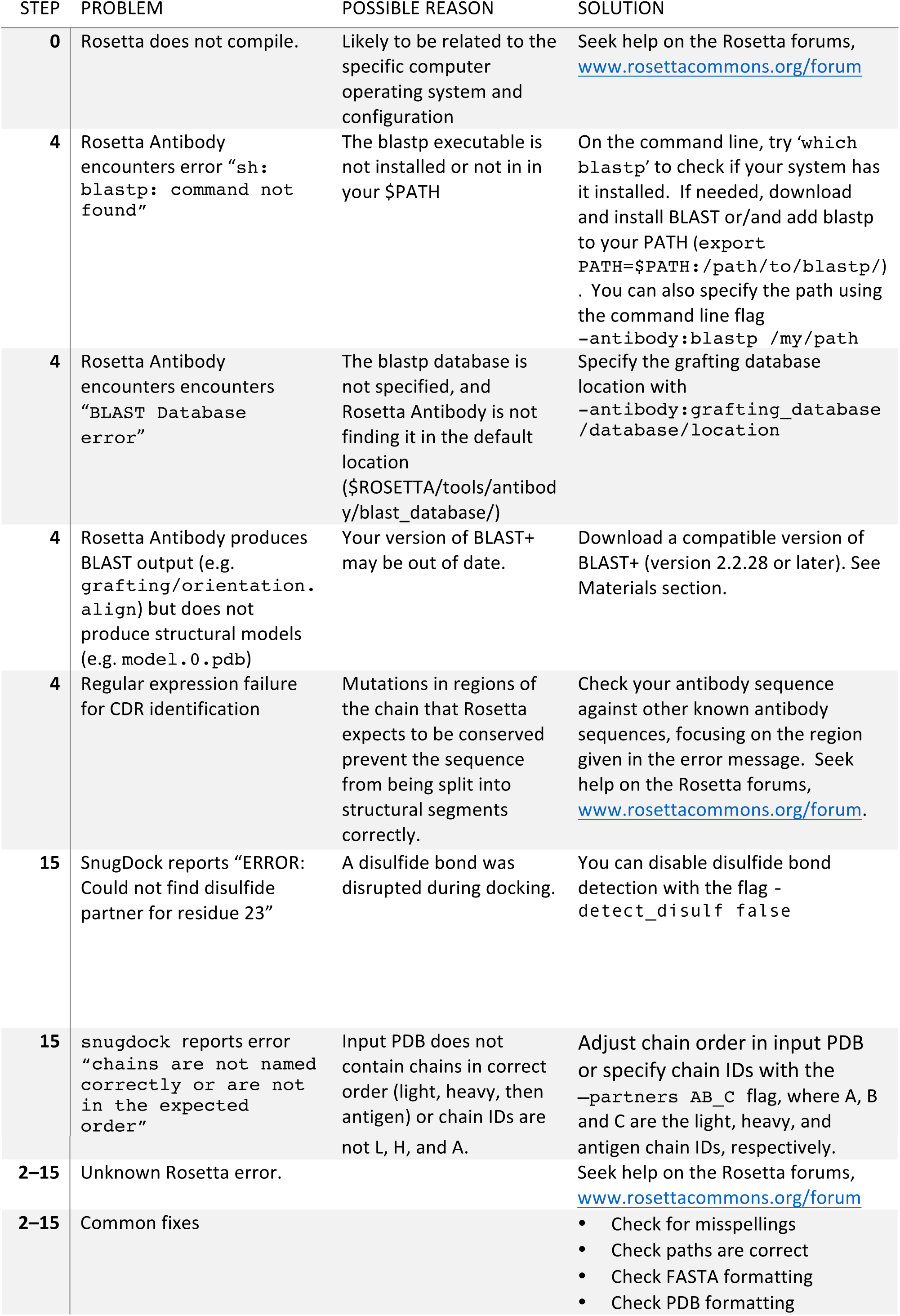

## ANTICIPATED RESULTS

The antibody structure prediction and docking methods described in this paper each produce a set of structural models that have been evaluated by a score function. In the case of antibody structure prediction, we have found through benchmarking and participation in the AMA that the accuracy of frameworks and non-H3 CDR loops can typically be expected to be within 1.0 Å RMSD of the coordinates in a crystal structure. When the model deviates more than 1.0 Å in RMSD from crystallographic coordinates it is usually because there is not a suitable known template in the PDB. These situations should become increasingly rare as more structures are deposited into the PDB, although heavily engineered antibodies should always be modeled with care.

The H3 loop accuracy is variable and depends both on length and V_L_–V_H_ orientation. Loop length is an important factor in the accuracy of *de novo* loop modeling methods because the search space increases exponentially with each additional residue in the loop. We expect accurate models of CDR H3 loops of length 14 or less^23^, but the top-scoring model may not be the most accurate. We therefore recommend using all ten models for downstream analysis. In AMA-II, we found that non-native V_L_–V_H_ orientations can lead to explicit interactions between the light chain and the CDR H3 loop that are indistinguishable from native interactions^28^. Using multiple V_L_–V_H_ orientation templates^18^ allows broader exploration of conformational space, sampling more low-scoring wells. Models generated from at least three different templates should be used to maximize the chance of capturing the native V_L_–V_H_ orientation.

Through benchmarking Ab–Ag docking, we have found that the accuracy of a complex model depends on the starting configuration of the partners and the accuracy of the models for each partner. SnugDock samples local conformation space, thus a good starting structure (within 8 Å) generally results in sampling a near-native conformation. Equally important is the quality of the initial unbound models; near-native models enable increased docking performance (see Table 1: B-B rigid body-docking vs. U-U rigid-body docking). We have found that docking a homology modeled antibody to the crystal structure of the unbound antigen typically results in at least one model of acceptable quality in the ten top-scoring models (Table 1).

## AUTHOR CONTRIBUTIONS

BDW, NM, SL, DK, JRJ, and JJG developed the current version of RosettaAntibody. SL developed ROSIE and implemented the RosettaAntibody and SnugDock server apps. BDW implemented SnugDock in Rosetta 3, JRJ benchmarked SnugDock’s performance. RF and NB wrote the procedure, codified the manual intervention steps developed by BDW, NM, and DK, and recorded timing information. BDW, NM, SL, JRJ, DK, RF, NB and JJG wrote the manuscript.

## ACKNOWLEDGMENTS

The authors wish to thank Arvind Sivasubramanian, Aroop Sircar, and Sidhartha Chaudhury for their development of the original RosettaAntibody, SnugDock, and EnsembleDock methods. Jianqing Xu refactored the antibody code. We also thank the members of the RosettaCommons for the continued development of the Rosetta Software Suite. ROSIE simulations are carried out, in part, within the Extreme Science and Engineering Discovery Environment (XSEDE), which is supported by National Science Foundation grant number ACI-1053575. BW, NM, JRJ and JJG are supported by National Institutes of Health Grant R01 GM078221. SL is supported by National Institutes of Health Grant R01 GM73151. DK is supported by the DARPA Antibody Technology Program (HR-0011-10-1-0052) and the Japan Society for the Promotion of Science (grant number 15H06606). RF is supported by the South-Eastern Norway Regional Health Authority (grant number 850703-6051-39788).

